# Attentional Networks during the Menstrual Cycle

**DOI:** 10.1101/717264

**Authors:** Zahira Z. Cohen, Neta Gotlieb, Offer Erez, Arnon Wiznitzer, Oded Arbel, Devorah Matas, Lee Koren, Avishai Henik

**Affiliations:** Department of Psychology and Zlotowski Center for Neuroscience, Ben-Gurion University of the Negev, Beer-Sheva, Israel; Department of Psychology, University of California Berkeley, California, United States; Soroka University Medical Center and School of Medicine, Ben-Gurion University of the Negev, Beer-Sheva, Israel; Rabin Medical Center and Sacker faculty of medicine, Tel-Aviv University, Tel-Aviv, Israel; Mindfulness Clinic, Beer-Sheva Mental Health Center, Beer-Sheva, Israel; Faculty of Life Sciences, Bar-Ilan University, Ramat-Gan, Israel

**Keywords:** Menstrual Cycle, Progesterone, Estradiol, Attention, ANT-I, ANT, Alertness, Arousal, Orienting, Executive

## Abstract

The menstrual cycle is characterized partially by fluctuations of the ovarian hormones estradiol (E2) and progesterone (P4), which are implicated in the regulation of cognition. Research on attention in the different stages of the menstrual cycle is sparse, and the three attentional networks (alerting, orienting and executive) and their interaction were not explored during the menstrual cycle. In the current study, we used the ANT-I (attentional network test – interactions) to examine two groups of women: naturally cycling (NC) – those with a regular menstrual cycle, and oral contraceptives (OC) – those using OC and characterized with low and steady ovarian hormone levels. We tested their performance at two time points that fit, in natural cycles, the early follicular phase and the early luteal phase. We found no differences in performance between NC and OC in low ovarian hormone states (Both phases for the OC group and early follicular phase for the NC group). However, the NC group in the early luteal phase exhibited the same pattern of responses for alerting and no-alerting conditions, resulting in a better conflict resolution (executive) when attention is oriented to the target. Results-driven exploratory regression analysis of E2 and P4 suggested that change in P4 from early follicular to early luteal phases was a mediator for the alerting effect found. In conclusion, the alerting state found with or without alertness manipulation suggests that there is a progesterone mediated activation of the alerting system during the mid-luteal phase.

---

Attention is a general cognitive process that can be subdivided into three networks; alerting, orienting, and executive (or cognitive control) (Petersen & Posner, 2012; Posner & Petersen, 1990). Alertness refers to the level of arousal, orienting refers to the spatial allocation of attention and executive refers to the process that maintains an appropriate problem solving set to attain a future goal (i.e., conflict resolution; Callejas, Lupiàñez, Funes, & Tudela, 2005; Callejas, Lupiáñez, & Tudela, 2004; Fan, McCandliss, Sommer, Raz, & Posner, 2002; Posner & Petersen, 1990; Welsh & Pennington, 1988).

Alertness can be either phasic or tonic; phasic alertness is a rapid change of attention following an external alerting event, and tonic alertness is an intrinsic state of arousal, that fluctuates on the order of minutes to hours (DeGutis & Van Vleet, 2010; Weinbach & Henik, 2013). The neural mechanisms supporting phasic alertness have shown to highly overlap with areas responsible for tonic alertness, including right inferior frontal, inferior parietal, and anterior cingulate brain regions (DeGutis & Van Vleet, 2010; Sturm & Willmes, 2001), modulated by noradrenaline (NE), that is secreted by the locus coeruleus (LC) (The LC-NE system; Aston-Jones & Cohen, 2005). Orienting of attention in space can be attained overtly through eyes or head shifts or covertly through attention movements. The orienting network is regulated by frontal and parietal brain regions such as the superior parietal lobe, temporal parietal junction and frontal eye field (Corbetta, Kincade, Ollinger, McAvoy, & Shulman, 2000; Corbetta & Shulman, 2002; Posner, 1980), and is modulated by dopamine and acetylcholine (Lundwall, Guo, & Dannemiller, 2012; Phillips, McAlonan, Robb, & Brown, 2000; Witte, Davidson, & Marrocco, 1997). The executive network involves frontal and prefrontal brain areas such as the anterior cingulate cortex (ACC) and often the lateral prefrontal cortex (Botvinick, Cohen, & Carter, 2004; Fan et al., 2002), and involves dopamine and serotonin (Rosario, Pozuelos, & Combita, 2015). According to Callejas et al., (2004), these three attention networks interact with each other.

Attentiveness varies according to intrinsic and extrinsic factors. During the menstrual cycle, attentiveness may be modulated by changes in ovarian hormones, such as estradiol (E2) and progesterone (P4). The concentrations of these hormones fluctuate throughout the menstrual cycle, and these hormonal changes are associated with alterations in cognitive functions, including attention.

During the early follicular phase of the menstrual cycle (i.e., the days after the first day of menstruation) E2 and P4 levels are low. As the ovarian follicles develop, E2 levels gradually rise, peaking prior to ovulation. In the luteal phase (i.e., the 14 days between ovulation and the following menses), E2 levels decrease whereas P4 secretion from the corpus luteum gradually rises to support implantation of a fertilized ovum. If pregnancy does not occur, P4 levels decline and menstruation takes place (Fillingim & Ness, 2000; Haimov-Kochman & Berger, 2014).

Menstrual cycle and ovarian hormones-related changes have been studied in the context of cognitive abilities (e.g., Sacher, Okon-Singer, & Villringer, 2013; Symonds, Gallagher, Thompson, & Young, 2004), and specifically sustained, divided and selected attention (Pletzer, Harris, & Ortner, 2017; Thimm, Weis, Hausmann, & Sturm, 2014), spatial attention (Brötzner, Klimesch, & Kerschbaum, 2015; Hausmann, 2005) and executive attention (conflict resolution) (Hatta & Nagaya, 2009). Beaudoin and Marrocco (2005) showed that menstrual cycle phase effects the spatial allocation of attention, orienting, and alertness (independently). Upadhayay and Guragain (2014) found that females in the luteal phase may have advantages in some executive control tasks (Stroop and go-no-go tasks, but not in the flanker task), compared to males. However, Solís-Ortiz and Corsi-Cabrera (2008) reported decreased performance in executive functions, measured by the Wisconsin Card Sorting Test, in luteal women and improved performance in sustained attention, measured by the Continuous Performance Test. Brötzner, Klimesch, and Kerschbaum (2015) reported that higher P4 was related to faster response in a visuospatial cued attention paradigm. However, other studies found no difference between luteal and follicular women in the Stroop task (Hatta & Nagaya, 2009), auditory divided attention (Mordecai, Rubin, & Maki, 2008) and sustained attention tasks (Matthews & Ryan, 1994). The discrepancy in the literature may originate in the different tasks, small sample sizes, between-subjects hormone level variability, time of day, the specific day within the luteal phase and interactions between attention networks that were not controlled. Conducting experiments using a within-participant design, as well as measuring the interactions between different attention components, may elucidate some of these discrepancies.

Hormonal contraceptives suppress ovarian hormone production via negative feedback on the hypothalamic-pituitary-gonadal (HPG) axis. As contraceptives inhibit the release of LH and FSH, follicular development is suppressed, and ovulation does not occur (Mishell, Kletzky, Brenner, Roy, & Nicoloff, 1977; Rivera, Yacobson, & Grimes, 1999). The HPG suppression in women taking hormonal contraceptives is associated with structural, physiological, and functional changes, which are related, directly or indirectly, to attention. Differences in brain structures have been reported in women using hormonal contraceptives, including higher volume of gray matter in prefrontal cortices, compared to naturally cycling women (Pletzer et al., 2010). Additionally, women with hormonal suppression exhibit altered blood-oxygen-level-dependent (BOLD) signal and regional cerebral blood flow (rCBF) responses to emotional stimuli, showing increased activity in the amygdala and decreased activity in prefrontal regions, with no behavioral differences. This alteration in BOLD signal was restored when E2 or P4 were administered (Berman et al., 1997; Gingnell et al., 2013). When comparing the resting states of women who are naturally cycling to those using hormonal contraceptives, marked differences in brain connectivity were found, particularly in the anterior cingulate cortex and left middle frontal gyrus, areas involved in cognitive and emotional processing (Petersen, Kilpatrick, Goharzad, & Cahill, 2014).

## The Current Study

The three attentional networks are influenced by changes in ovarian hormones and these networks interact with each other. Studying networks’ interactions across different stages of the menstrual cycle can advance the understanding of attention and its regulation in women. According to our knowledge, no study has tested the influence if the menstrual cycle of the interactions of the three attentional networks using direct measures of ovarian hormones. In order to do so, we used the attentional networks test – interaction (ANT-I; Callejas et al., 2004), which allows independent manipulation of three attention networks; alertness, orienting and executive, and measures their interactions with one another (see Methods section 2.4.3 for an elaboration of the specific pattern of interactions). We measured the performance on the ANT-I among two groups of women: naturally cycling (NC) women and oral contraceptives (OC) women. The NC women do not use hormonal contraceptives and are assumed to have a regular menstrual cycle with natural hormone fluctuations. The OC women use contraceptives that maintain low steady hormonal levels throughout the month. Each participant was measured during two sessions in different stages of the menstrual cycle, the early follicular phase (when E2 and P4 levels are low) and early luteal phase (when E2 and P4 levels are relatively high) (See Methods section 2.3 for elaboration).

The OC group differs from the NC group in more than one aspect, hence cannot serve as a direct control group without exposing the analysis to confounds. Therefore, we had two hypotheses, one for each group: 1) For the OC group, there would be no behavioral change between the early follicular and early luteal phases in attentional networks performance, reflecting the similar (low) hormonal levels in these two time points. 2) For the NC group, we expected to find a difference between the early follicular and early luteal phases, reflecting different hormonal levels between the two time points. Specifically, we expected to find an alteration in the interaction between the three attentional networks.

## Materials and Methods

### Subjects

Total of 71 female right-handed students participated in this experiment in two recruitment phases. See Table 1 for the distribution of the number of women in each group and order. Out of this group, 45 participants met the inclusion criteria and were included in the analysis (see “exclusion criteria” section). The mean age of the OC group was 23.4± 1.3 and the NC group 22.9 ± 2.1 years. The range was 19-27 years. Participants were paid for their participation or received course credit for “Introduction to Psychology” course. None of the women used any neuroactive substance or reported on a diagnosed mental disorder.

**Table 1.** Participants’ allocation for order condition (early follicular first / early luteal first) in each group (OC / NC). In parentheses, the number of participants prior exclusions, and out of parentheses, the final number of participants that were included in the analyses.

| Group | OC | NC | Total |
| --- | --- | --- | --- |
| Order |  |  |  |
| Early follicular first | 14 (19) | 11 (17) | 25 (36) |
| Early luteal first | 10 (17) | 10 (18) | 20 (35) |
| Total | 24 (36) | 21 (35) | N = 45(71) |

### Exclusion criteria

Participants were screened using the Ben-Gurion University’s recruiting questionnaire for having a history of regular (i.e., 27 to 29 day) menstrual cycles, with no history of skipping cycles. For the NC group, we recruited participants with at least six months of no contraceptive use prior to the time of study, and for the OC group we recruited participants with at least six months of using the same oral hormonal contraceptive (i.e., birth control pills), that contained both E2 and P4 in a constant ratio throughout the month. 71 participants performed the experiment. Post-hoc analysis of their actual cycle revealed a median cycle length of 29 days. 26 participants were excluded from the analysis for the following reasons^1^: 1) self-reported scores in the questionnaires of both sessions indicated severe depression, severe anxiety or premenstrual dysphoric disorder (PMDD) (n = 12, OC: 4, NC: 8); 2) the actual menstrual cycle, that was measured following the experiment, exceeded 31 days (n = 12, OC: 3, NC: 9); 3) technical problems with running the experiments (n = 4, OC: 2, NC: 3); or 4) participants changed their pill administration (n = 3, OC: 3).

### Procedure

According to the reported length of the menstrual cycle, participants were scheduled for 2 sessions; one session in the early follicular phase (i.e., for 28 length cycle – the 4^th^ day, and for 27 length cycle – the 3^rd^ day) and the other was in the early luteal phase (i.e., for 28 length cycle – the 18^th^ day, and for 27 length cycle – the 17^th^ day). We chose the 4^th^ and 18^th^ cycle days in order to 1) hit the early follicular phase (when E2 and P4 levels are low) and early luteal phase (when E2 and P4 levels are relatively high), and 2) optimizing the difference between the two sessions to about 14 days (in a 28^2^ days of menstrual cycle). Starting session was counterbalanced, so that half of the participants started in the early follicular session, and half started in the early luteal session.

We collected the data during two consecutive recruitment phases. The first recruitment phase included the ANT-I and saliva collection for E2 and P4 measurement, and the second recruitment phase included only a replication of the behavioral ANT-I with different participants. No statistical difference was found between the two cohorts of the study^3^. Hence, we report the behavioral results of the two recruitment phases as one data set.

Before the first session participants were instructed regarding saliva collection for E2 and P4 including (a) going to sleep and waking up at the same hour for both sessions; (b) refraining from eating or drinking (except water) for an hour before the experiment; (c) refraining from brushing teeth prior to saliva collection to avoid bleeding gums.

At the beginning of each session, participants sat in front of a computer, in an isolated, lit experimental room and completed a questionnaire that determined (a) personal and academic information; (b) the current day in the menstrual cycle; (b) premenstrual symptoms, mood and affect related questionnaires; (c) and awakening time. Later, the participants gave a saliva sample, which was immediately frozen at −20°C and started the ANT-I task.

Participants were instructed to inform us when they received their next menses to estimate their time of ovulation and update the information about the chosen session days.

### Materials

#### Questionnaires

*The Beck Anxiety Inventory* (BAI; Beck, Brown, Epstein, & Steer, 1988). The BAI is a self-report measure designed to assess anxiety symptoms. Participants are asked to rate how much they have been bothered by each of the 21 anxiety related symptoms over the past week on a 4-point scale, ranging from 0 to 3. Scoring above 21 points (moderate anxiety) in both phases was the exclusion criterion.

*The Beck Depression Inventory II* (BDI-II; Beck, Steer, & Brown, 1996). The BDI-II is inventory designed to assess current severity of depression. Participants are asked to rate the severity of each of the 21 depression related symptoms and attitudes on a 4-point scale, ranging from 0 to 3. Scoring above 21 points (moderate depression) in both phases was the exclusion criterion.

*The premenstrual symptoms screening tool for clinicians* (*PSST*; Steiner, Macdougall, & Brown, 2003). The PSST is a 19-item instrument consisting of two domains: the first domain includes 14 items related to psychological, physical, and behavioral symptoms and the second domain evaluates the impact of symptoms on functioning within five life aspects (a-e). Each item is rated on a four-point scale ranging from 0 to 3. For a diagnosis of PMDD, the following criteria must be present: (1) at least one of the symptoms (1 to 4) are scored 3; (2) in addition, at least four of the symptoms (1 to 14) are scored 2 or 3; and (3) at least one of a, b, c, d, or e are scored 3. The exclusion criterion for our study is a PMDD diagnosis.

### Quantitation of estradiol and progesterone

On the day of assaying, saliva samples were thawed and centrifuged for 15 min at 3000 rpm. E2 and P4 were quantitated in saliva extracts using commercial enzyme-linked immunosorbent assays (ELISA; Salimetrics, Ann Arbor, MI, USA: progesterone 1-1502; estradiol 1-3702) according to the manufacturer’s recommendations. Intra-assay repeatability was determined using three duplicates of a pool (n = 6) on the same ELISA plate for all hormones. The calculated coefficient of variation for progesterone was 11.7% and for estradiol 5.35%. Inter-assay repeatability using 2 duplicates (n = 4) was 10.39% for progesterone and 10.92% for estradiol.

### ANT-I

The ANT-I resembled the task used at Callejas et al. (2004). 1) Phasic alertness was manipulated by administrating an alerting signal before a task. The alerting effect refers to the difference in reaction time (RT) between alerting condition and the no-alerting condition. 2) Changes in the orienting network were achieved by using valid or invalid cues before a target stimulus. A valid cue is expected to create a faster response than an invalid cue (i.e., presenting a cue on the opposite side of the target). The validity effect refers to the difference in RT between the valid and invalid conditions. 3) Executive network (cognitive control) was studied by using a stimulus that one aspect of it needs to be focused and another aspect of it needs to be ignored. Conflict resolution requires time; therefore in a conflicted trial, RT is slower than in non-conflicted trials (Callejas et al., 2005, 2004; Fan et al., 2002; Posner & Petersen, 1990). Congruency effect refers to the difference in RT between congruent and incongruent conditions. The interactions of each network with the others are as follows: a) The alerting network spatially influences the executive network, creating a larger congruency effect on alerting condition, compared to no-alerting condition, b) The orienting network influences positively on the executive network, causing a smaller congruency effect for valid (compared to invalid) condition, and c) The alerting network causing a faster orienting, creating a bigger validity effect for alerting (compared to no-alerting) condition. The interaction between the three attentional networks was not analyzed or discussed in the study of Callejas et al., but could be extracted from the reported data; the alerting network influence executive network differently, when activating the orienting network. Specifically, for the alerting condition, when observing valid condition, the congruency effect was smaller (vs. invalid condition). However, for the no-alerting condition, the congruency effect was about the same for valid and invalid conditions.

Each trial began with a fixation point presented for a duration ranging between 400 milliseconds (ms) and 1,600 ms. In half of the trials, a 50 ms alerting signal (2,000 Hz) was presented along with the fixation point. After a 400 ms stimulus onset asynchrony (SOA), in 2/3 of the trials, an orienting cue was presented for 50 ms below or above the fixation point: 1/3 of the trials the cue was valid, and 1/3 of the trials the cue was invalid. In the other 1/3 of the trials, only fixation point was presented, with no orienting cue (i.e., no cue). After another 50 ms SOA, a target arrow and surrounding flankers were presented below or above the fixation point. Participants were instructed to focus their attention on the middle arrow and ignore the surrounding flanking arrows. The target arrow and four flankers were presented either in congruent direction (i.e., pointing in the same direction) or incongruent direction (i.e., the target arrow was pointing in the flankers’ opposite direction), for 3,000 ms or until a response. Each trial sequence lasted for a total time of 4,450 ms, which means that after the response, the fixation point remained on screen for a varying amount of time. See Figure 1 for trial sequence example.

**Fig. 1.**
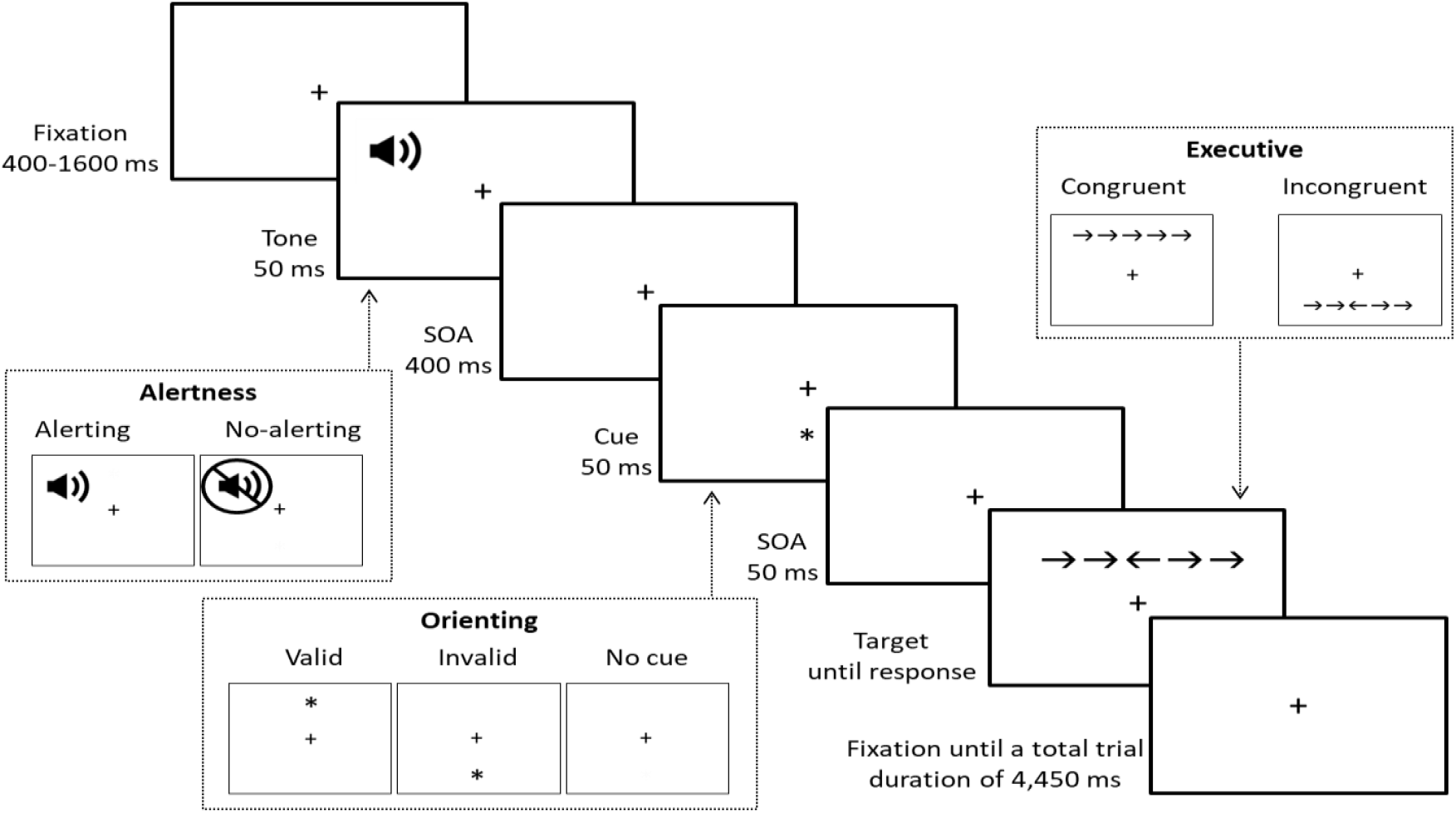
Schematic trial of the ANT-I with an example of alerting trial, invalid spatial cue, and incongruent target. Participant needs to ignore the flanking arrows to the right side and respond with the left key since the middle arrow is pointing left.

Each session started with a practice block of 10 accuracy feedback trials. After the practice block there were 6 blocks of 48 trials each (2 [alerting/no-alerting] X 3 [valid/invalid/no cue] X 2 [congruent/incongruent] X 4 repetitions), for a total of 288 trials altogether.

### Analysis

To test the differences between groups and time in the ANT-I, a 6-way mixed ANOVA was conducted. The between-participants variables were 1) group (NC / OC); 2) order (early follicular first / early luteal first) and the within-participants variables; 3) time (early follicular / early luteal), and the three ANT-I variables; 4) congruency (congruent / incongruent); 5) validity (valid / invalid; excluding no cue trials); and 6) alertness (alerting / no-alerting).

To further explore the behavioral results, we tested E2 and P4 changes from the early follicular to the early luteal phase of the first recruitment phase (N = 20; OC: 10, NC: 10). Since ANCOVA is not the most suitable analysis for within-participant variables, we created a new variable, RT interaction, and conducted a simple regression analysis. RT interaction was the dependent variable that represents the main behavioral finding–the simple two-way interaction of validity and congruency within each level of alertness. It is calculated, for each participant, within each level of alertness, as the congruency effect of the valid condition subtracted from the congruency effect of the invalid condition (i.e., invalid [incongruent – congruent] – valid [incongruent – congruent]). RT interaction = 0 represents the main effects of validity and congruency only, with no interaction. As the RT interaction increases (or decreases) the interaction effect increase. This means that the two-way interaction is more evident. The independent variables for the RT interaction analyses were delta P4 and delta E2, which is the difference between the two sessions, within-participant. A value of 0 represents no difference between early luteal and early follicular phases, while a positive value represents higher hormone levels in the early luteal phase and a negative value represents lower hormone levels in the early luteal phase. To test the general differences between groups in age, depression, anxiety, P4 and E2, ANOVAs were conducted for each variable and between early follicular and early luteal phases.

## Results

### ANT-I

#### Preliminary analysis

As in Callejas et al. (2004), we included only correct trials, in the range of 200 ms and 1200 ms, which were 97% of the trials. First, we report the three-way interaction of the ANT-I (beyond time, group and order). The results replicated the observed findings, *F*(1, 41) = 14.3, *MSE* = 464, *p* = .01, *η*^2^_*p*_ = .26. That is, the alerting network influenced executive network differently, when activating the orienting network. Further analysis, to explore the pattern of differences in each alertness level, revealed that for the alerting condition the orienting X executive contrast was significant, *t*(41) = 5.89, *p* = .000001, *η*^2^_*p*_ = .46. Pattern of RT reveals that the congruency effect was smaller in the valid than the invalid condition. In contrast, for the no-alerting condition, the orienting X executive contrast was not significant, *t*(41) = 1.9, *p* = .064. That is, the congruency effect was about the same for valid and invalid conditions.

#### Main results - hypotheses testing

Since there was no main effect of order, F(1, 41) = 1.4, *p* = .21, and the highest interaction found was the 5-way interaction of time X group X congruency X validity X alertness, *F*(1, 41) = 6.7, *MSE* = 25,804, *p* = .01, *η*^2^_*p*_ = .14, we continued to analyze the results, based on our hypotheses. To test whether the ANT-I interaction differed between the two time phases, we carried out an interaction between comparisons contrast (i.e., mean interaction contrast). That is, we compared RT difference from the early follicular and the early luteal phases, between the three attentional networks: arousal, orienting and executive.

For the OC group (hypothesis 1), we found no significant difference in the contrast of time and the ANT-I variables, *t*(41) = 1.31, *p* = .2, *η*^2^_*p*_ = .04. This contrast indicates that the ANT-I interaction was not different between the two time phases. For the NC group (hypothesis 2), there was a significant difference in the contrast of time and the ANT-I variables, *t*(41) = 2.33, *p* = .02, *η*^2^_*p*_ = .11. This contrast indicates that there was a difference between the early follicular to the early luteal phases in the ANT-I pattern of response.

#### Additional analyses

In order to describe the pattern of the ANT-I, further analyses were done within each alertness level to explored the difference in the executive and orienting interaction. Following the main results, the analyses were done, separately, beyond time variable in the OC group, and within each of the time condition in the NC group (see Figure 2 for the RT pattern of results).

**Fig. 2.**
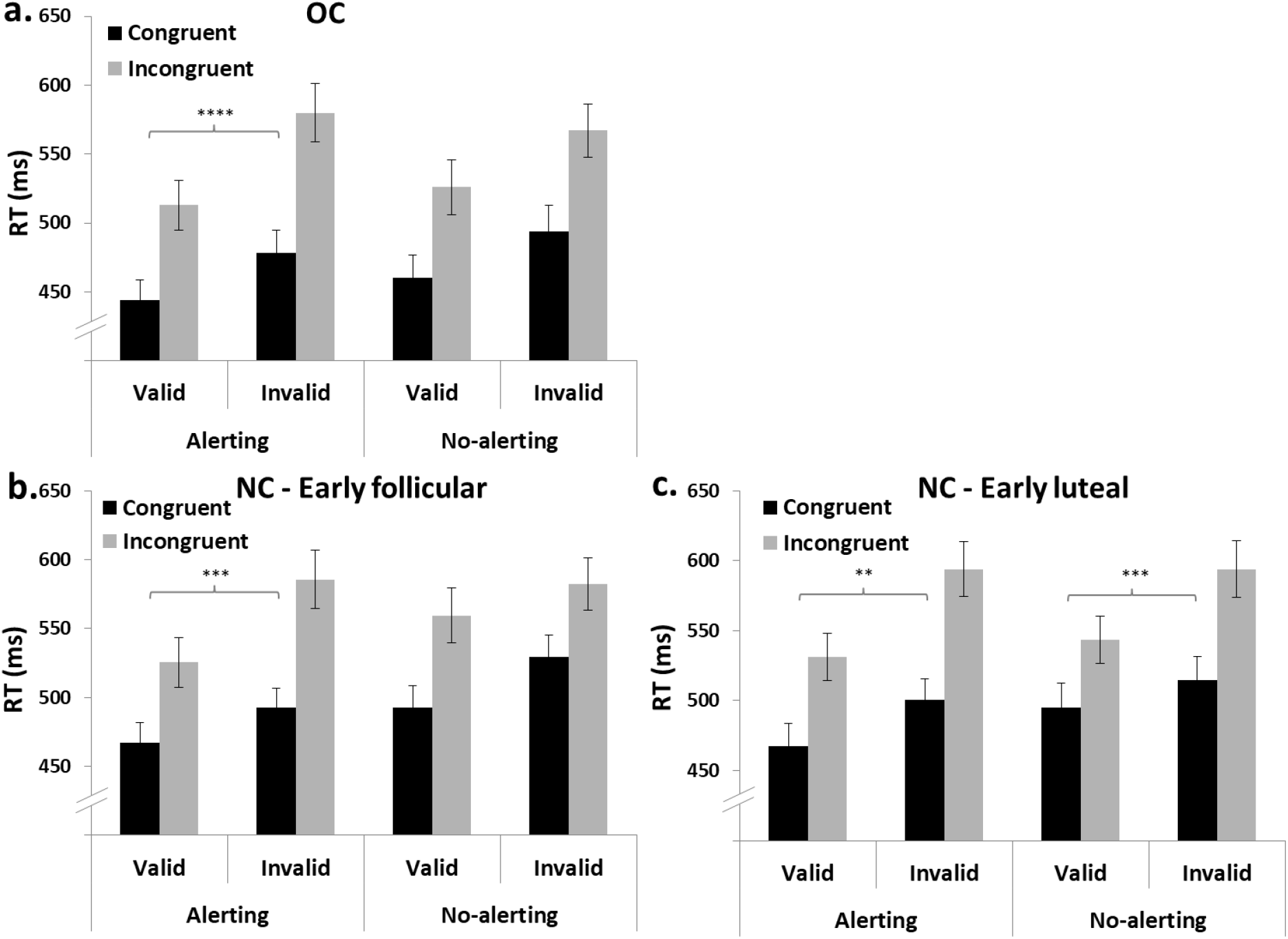
Response time (RT in ms) of OC group (a) and NC group – early follicular (b) and early luteal (c) phases, of congruent vs. incongruent, valid vs. invalid and alerting vs. no-alerting conditions. ** - *p* < .01, *** - *p* < .001, **** - *p* < .0001

For the OC group, the Bonferroni-corrected alpha was .0125. The ANT-I contrast was significant, *t*(41) = 2.88, *p* = .006, *η*^2^_*p*_ = .17. The RT pattern of the orienting X executive contrast in each alertness level resembled the main findings, presented in the preliminary analysis (alerting: *t*(41) = 4.38, *p* = .0001, *η*^2^_*p*_ = .32, no-alerting: *t*(41) = 1.29, *p* = .2, see Figure 2a).

For the NC group, the Bonferroni-corrected alpha was .00625. In the early follicular phase, the ANT-I contrast was significant, *t*(41) = 3.49, *p* = .001, *η*^2^_*p*_ = .21. The RT pattern the of orienting X executive contrast in each alertness level resembled the main findings, presented in the preliminary analysis (alerting: *t*(41) = 3.18, *p* = .001, *η*^2^_*p*_ = .21, no-alerting: *t*(41) = 1.5, *p* = .12, see Figure 2b). However, in the early luteal phase the ANT-I contrast was not significant, *t*(41) = 0.15, *p* = .88, *η*^2^_*p*_ = .0005 (see Figure 2c). Although this contrast was not significant, we further explored the orienting X executive contrast within each alertness level. By doing so, we could reveal the core differences in the ANT-I that are attributed to the menstrual phase. In the alerting condition (Figure 2c left), the orienting X executive contrast was significant, *t*(41) = 2.32, *p* = .002, *η*^2^_*p*_ = .21. Importantly, in the no-alerting condition orienting and executive contrast was significant as well (Figure 2c right), *t*(41) = 2.9, *p* = .006 *η*^2^_*p*_ = .17. In the early luteal phase, the no-alerting condition pattern of response was the same as in the alerting condition, resulting in a smaller congruency effect for valid trials than for invalid trials.

We conducted post-hoc analyses, comparing the ANT-I contrast between groups within each of the time phases. In the early follicular phase, there was no difference between groups in the ANT-I contrast, *t*(41) = 1.9, *p* = .07, *η*^2^_*p*_ = .08. In contrast, in the early luteal phase, this difference was significant, *t*(41) = 2.02, *p* = .049, *η*^2^_*p*_ = .09. These analyses showed that the pattern of interaction in the ANT-I is similar between groups in the early follicular phase but different in the early luteal phase, strengthening our within-participant analysis.

### Delta P4 and Delta E2 effects on RT interaction

Behavioral-results-driven regression analysis was conducted in order to explore the effects of delta P4 and delta E2 on the RT interaction found in the early luteal phase (See Figure 3 for the specific effect of RT interaction and group in the alerting and no-alerting conditions). Corresponding with the behavioral ANT-I analysis, in the OC group the RT interaction effect is evident only in the alerting condition, creating a difference between alerting and no-alerting condition, *t*(18) = 2.15, *p* = .04, *η*^2^_*p*_ = .21, while in the NC group the RT interaction effect is evident in both alerting and no-alerting condition, with no difference between the two conditions, *t*(18) = .47, *p* = .62, *η*^2^_*p*_ = .01. Meaning, the simple interaction effect of validity and congruency was evident in alerting condition (of both groups) and in no-alerting condition only for the NC group.

**Fig. 3.**
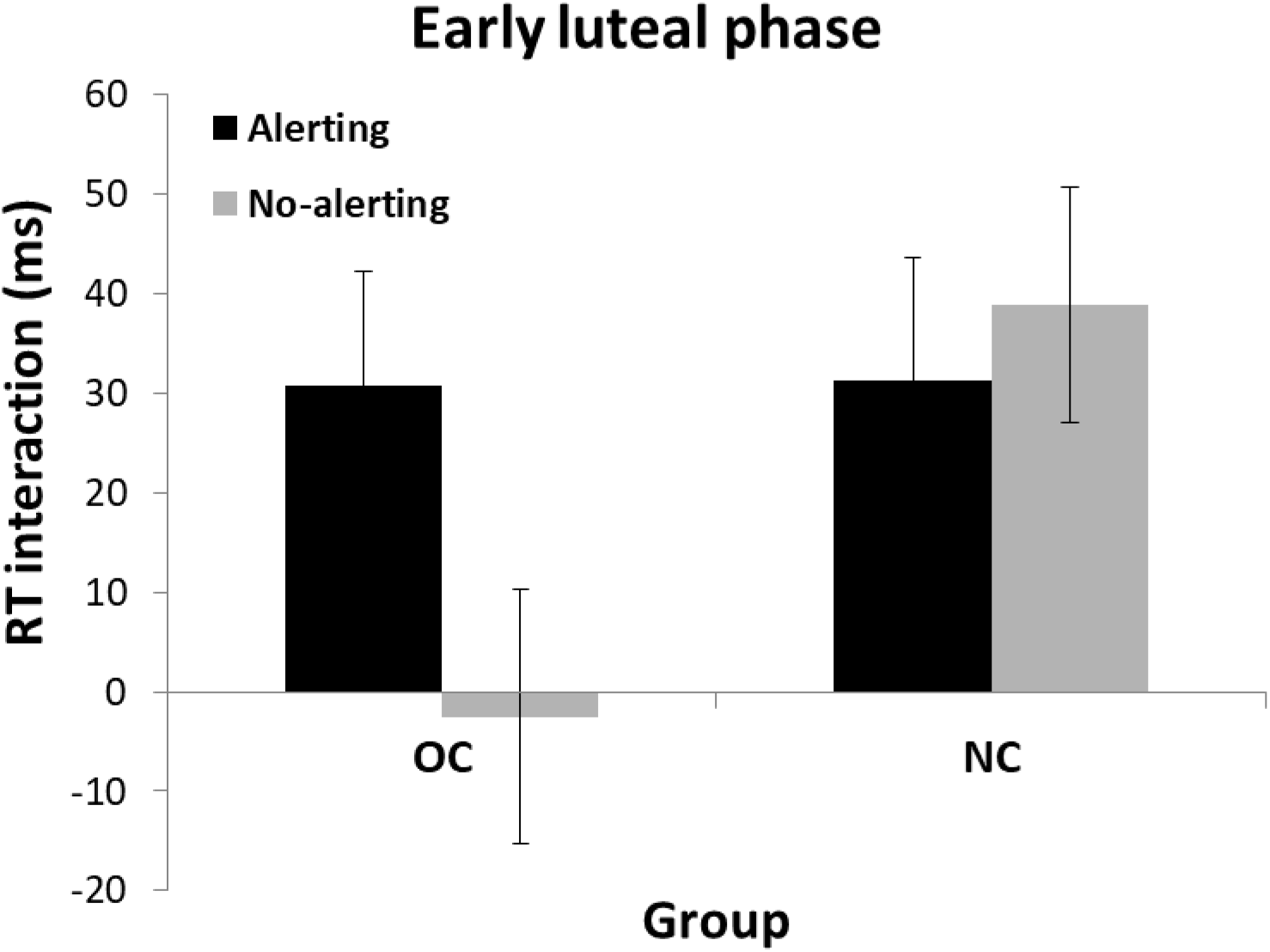
RT interaction (i.e., invalid [incongruent – congruent] – valid [incongruent – congruent]) in the early luteal phase within each of the alerting conditions (alerting / no alerting) between groups (OC / NC): RT interaction = 0 main effects of validity and congruency only, with no interaction. As the RT interaction increases (in absolute value) the interaction effect increases.

Accordingly, we used RT interaction as a dependent variable and delta E2 and P4 as an independent variable in separate regression analyses. We found that in the alerting condition of the early luteal phase, there was no linear relationship between the RT interaction and delta P4, *r* = .162, *p* = .248 (see Figure 4 higher panel for delta P4 scatter plot), or between RT interaction and delta E2, *r* = .021, *p* = .466. Consistent with our behavioral results, the difference between early follicular and early luteal levels of P4 or E2 was not related to the RT interaction of alerting condition. However, in the no-alerting condition, regression analysis showed that there was a correlation between the RT interaction and delta P4, *r* = .496, *p* = .013 (see Figure 4 lower panel for delta P4 scatter plot), but not in the RT interaction and delta E2, *r* = -.1, *p* = .342.

**Fig. 4.**
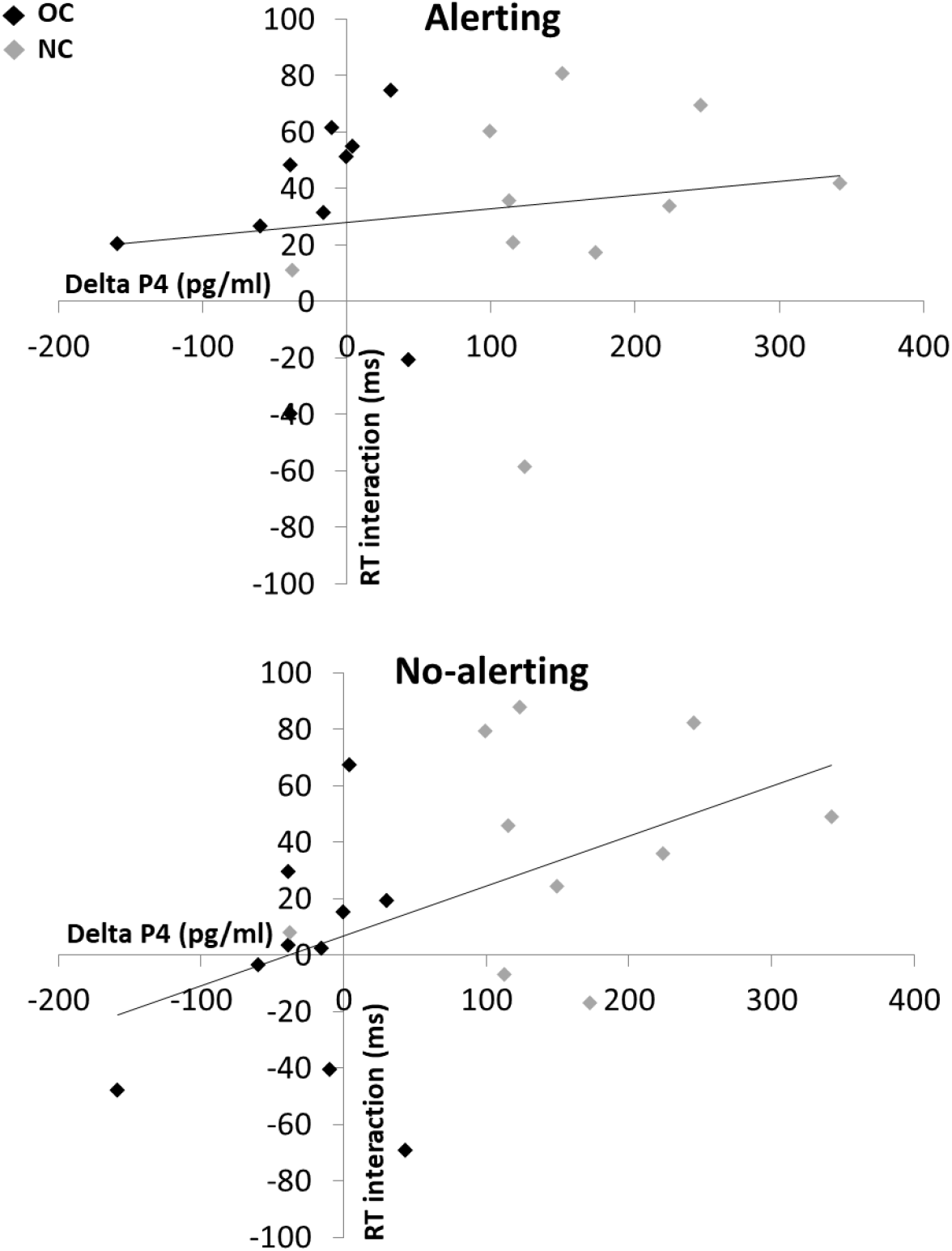
Mean RT interaction in the early luteal phase (i.e., invalid [incongruent – congruent] – valid [incongruent – congruent]) and delta P4 (P4 in the early follicular phase – P4 in the early luteal phase) for the NC group (grey) and the OC group (black). In the alerting condition, RT interaction is not associated with the levels of delta P4; In contrast, in the no-alerting condition, RT interaction is associated with the levels of delta P4; higher levels of delta P4 are associated with higher RT interaction.

Accordingly, we investigated whether delta P4 mediated the effect of group (0 = OC, 1 = NC) on RT interaction. Step 1 results of the mediation model indicated that when ignoring the mediator - delta P4, group was a significant predictor of RT interaction, *b* = 41.435, *β* = .49, *SE* = 17.397, *p* = .028. Step 2 showed that the mediator, delta P4, was a significant predictor to RT interaction, *b* = .177, *β* = .496, *SE* = .073, *p* = .026. Step 3 showed that group was a significant predictor of delta P4, *b* = 178.856, *β* = .753, *SE* = 36.799, *p* = .0001. Lastly, step 4 showed that group was no longer a significant predictor of RT interaction after controlling for the mediator, delta P4, *b* = 22.729, *β* = .269, *SE* = 26.547, *ns*. Approximately 28% of the variance in RT interaction was accounted for by the two predictors (*R*^2^ = .277). A Sobel test was conducted and confirmed the mediation path (*z* = 2.169, *p* = .03). These results suggest a mediation of delta P4 on group and RT interaction (see figure 5 for the mediation model).

**Figure 5.**
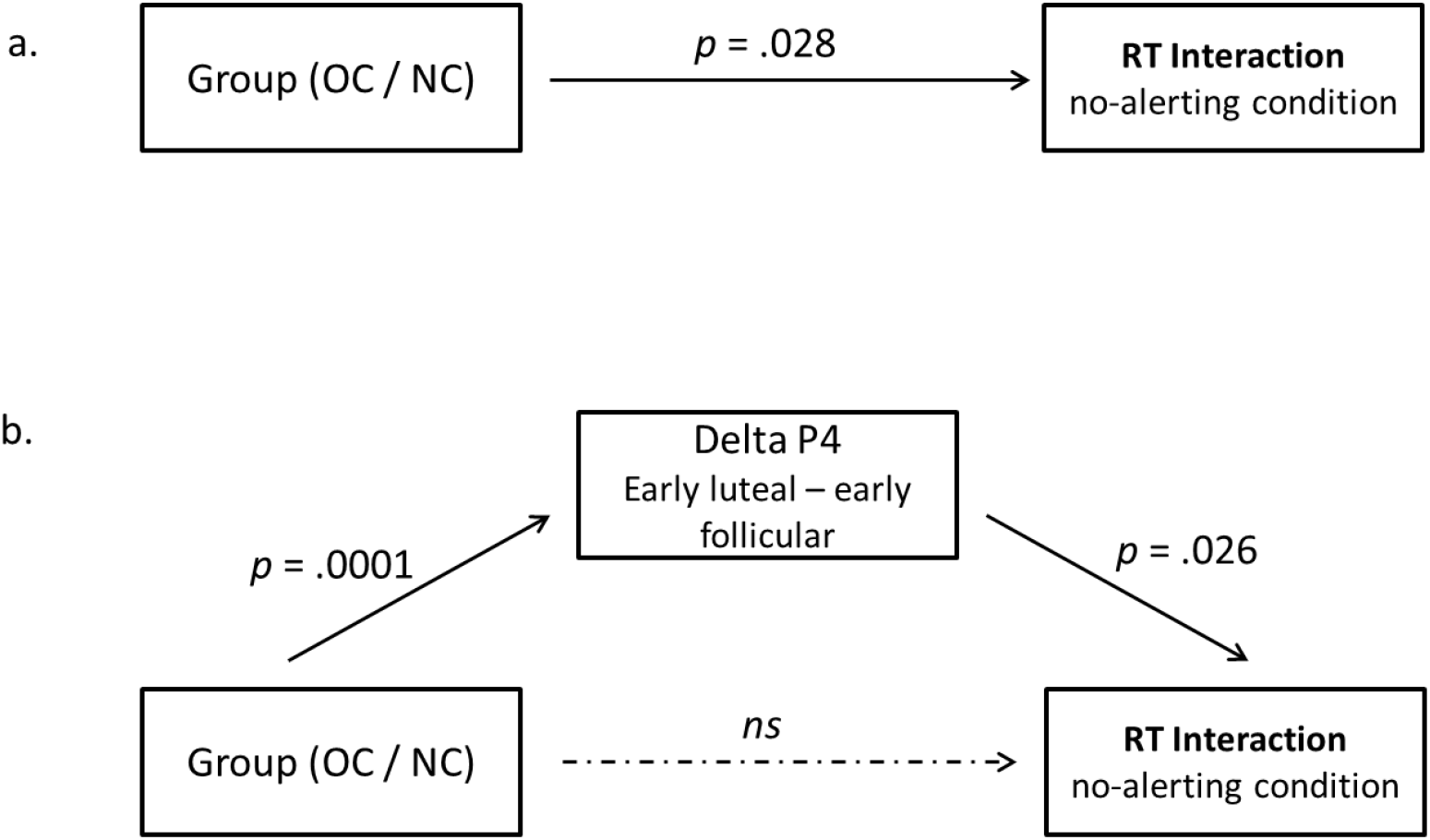
a. Group significantly predicts RT interaction (total effect; step 1). b. Direct effect and indirect effect mediated through delta P4 (steps 2-4).

Behavioral-results-driven regression analysis suggests that higher levels of P4, as in the early luteal phase among the NC group, are associated with increased alertness in the no-alerting condition, resulting in an interaction between orienting and executive networks.

### General group differences

No significant differences were found between the early follicular and the early luteal phases between OC and NC groups in age and depressed mood. The only significant difference between groups and time was in anxiety levels measured by the BAI. The two-way interaction of time and group was significant, *F*(1, 43) = 12.93, *MSE* = 13.05, *p* = .001 *η*^2^_*p*_ = .23. Simple effect of time within each group revealed that the levels of self-reported anxiety among the NC group were higher in the early follicular phase (vs. early luteal phase; *p* = .05), while in the OC group, the levels of self-reported anxiety were higher in the early luteal phase (18^th^ day) (vs. 4^th^ day; *p* = .01). The levels of anxiety between groups were not different in the early luteal phase, but only in the early follicular phase (p = .0001). Nevertheless, none of the participants met the cut-off for severe anxiety in both phases (NC mean (SD); early follicular: 12.57 (7.14), early luteal: 9.95 (5.8), OC; early follicular: 4.41 (4.18), early luteal: 7.29 (5)).

Salivary P4 ANOVA between groups (OC / NC) within time (early follicular / early luteal) demonstrated a significant two way interaction, *F*(1, 18) = 23.623, *MSE* = 3,385, *p* = .0001, *η*^2^_*p*_ = .56. Women from the NC group in the early luteal phase had significantly higher P4 levels (M = 261.03, SD = 26.03) than in early follicular phase (M = 106.42, SD = 23.35), t(18) = 5.94, p = .00001, *η*^2^_*p*_ = .66, while women from the OC group had no significant difference between early luteal (M = 83.25, SD = 26.03) and early follicular phase (M = 107.51, SD = 23.35), t(20) = −0.61, p = .54, *η*^2^_*p*_ = .01.

Salivary E2 ANOVA between groups (OC / NC) within time (early follicular / early luteal) showed no significant two way interaction, *F*(1, 18) = 1.37, *MSE* = .246, *p* = .257, nor main effect of group, *F* < 1, or time, *F*(1, 18) = 1.24, *MSE* = .246, *p* = .279 (NC: M = 2.3335, SD = .28, OC: M = 2.0007, SD =.295).

## Discussion

In this study, we examined the effects of menstrual cycle on the three attentional networks. Specifically, we tested hormonal changes between the early follicular and early luteal phases and their interactions with alerting, orienting, and executive networks. The behavioral results show that in the OC group—women that use oral contraceptives and have low and steady hormones levels—there was no significant difference between the two time points. In contrast, in the NC group—women with a natural menstrual cycle that are not using hormonal contraceptives showed a difference between early follicular and early luteal phases. In the early follicular phase, the response pattern did not differ from OC, replicating main effects and interactions of the ANT-I (that will be elaborated below). However, in the early luteal phase, the alerting system was activated, even when there was no alerting signal. This effect resulted in a smaller congruency effect for valid trials (vs. invalid trials). Results-driven regression analysis suggested that the change in progesterone levels from early follicular to early luteal phases in NC group is a mediator of the behavioral effect found.

The findings reported herein are in agreement with previous reports by Callejas et al. (2004; 2005), indicating that the three attentional networks modulate each other. This was evident as an overall effect (beyond group), in both phases among the OC group and in the early follicular phase among the NC group (in which E2 and P4 levels are low and about equal to the levels of the OC groups). The influence of the three attentional networks on each other was as follows: alerting network influenced the executive network differently when responding was to the oriented vs. the non-oriented location. When alerted, the ability of resolve a conflict (i.e., the congruency effect) was stronger (i.e., the difference between incongruent and congruent was smaller) when the attention was oriented to the same place as the conflicted stimuli (i.e., for valid trials, compared to invalid trials). When non-alerted, the ability to resolve conflict was not different for oriented and non-oriented location.

Although studies among women using hormonal contraceptives show alterations in brain structure, function, functional connectivity and resting state (Griksiene & Ruksenas, 2011; Mordecai et al., 2008; Petersen et al., 2014; Pletzer et al., 2010), we did not find any behavioral difference between the early follicular and the early luteal phases among the OC group. It might be that, as in Berman et al. (1997), there is a dissociation between the behavioral and neural manifestation of these differences. In the behavioral manifestation, the OC group showed the same pattern or responses and was found to be a good control for the effects of menstrual cycle on the attentional networks. This suggestion is strengthened by the fact that we did not find a difference between OC and NC groups in the early follicular phase but did find a difference in the early luteal phase. If we were to find a difference between groups in the early follicular phase (in which both groups are not under the immediate effect of using contraceptive) they could have been attributed it to the structural, more profound, difference between groups (i.e., brain volume etc.) or to other (not measured) differences between groups. Thus, the difference we report here of the NC group in the early luteal phase can be attributed to changes related to P4 and E2 levels (that did not occur in the OC group). Nonetheless, the analysis of the ANT-I and hormones levels were done, *a priori*, in two time points among each group, so that we could discuss the effect of menstrual cycle separately from the possible effects of hormonal contraceptives.

Our study is the first to examine the modulating effect of P4 on the three attentional networks and to find a P4-associated alerting state: Meaning, higher levels of P4 were associated with the alertness-like effect found in the no-alerting condition. Specifically, among the NC women, we found that changes in P4 levels (low in the early follicular phase and high in the early luteal phase) mediated the behavioral effect found in the early luteal phase. When P4 levels were higher, the ability to resolve a conflict (i.e., the congruency effect) was stronger (i.e., the difference between conflict and non-conflict trials are smaller) when the attention was allocated to the same location as the conflict stimuli (i.e., for valid trials, compared to invalid trials). This profound effect was found in both alerting and no-alerting conditions, suggesting that P4 may induce an alerting/arousal state.

Studies show that P4 was found to increase tonic inhibition of networks processing irrelevant information, improving cognitive processing, and specifically spatial attention (e.g., Brötzner et al., 2015). Alerting state broadens the attentional spotlight and increases the accessibility of salient visuospatial cues (Weinbach & Henik, 2013). We might carefully suggest, from an evolutionary perspective, that a higher alerting state during the early luteal phase (when P4 levels are higher) is advantageous in contributing to the safety of a potential pregnancy, serving as protective awareness from unexpected threats (See also Brötzner et al., 2015).

The relation between P4 and alertness may originate in the LC, the source of the brain’s NE (Aston-Jones & Cohen, 2005). Studies in several species, including humans, have shown that P4 and E2 receptors are expressed in the noradrenergic neurons in the LC and fluctuate in relation with hormonal event across the estrous cycle (in mice and rats), and that P4 stimulates the activity of LC noradrenergic neurons (Alonso-Solís et al., 1996; Genazzani et al., 2000; Helena et al., 2009; Helena et al., 2006; Scott et al., 2000; Szawka, Rodovalho, Monteiro, Carrer, & Anselmo-Franci, 2009). Although neural *de novo* synthesis of P4 may also play a role in stimulating the LC, it is feasible that circulating P4 in the early luteal phase contribute to an increased state of tonic alertness/arousal.

The inverted-U relationship between LC activity and attention tasks, demonstrated in Aston-Jones and Cohen (2005), suggests that there is an optimal level of LC activity. Moderate LC tonic activity and prominent phasic LC activity (following an alerting cue) is the optimal level, while low and high LC activity can impair attention and performance. To monitor for task-related utility, the LC is connected to the ACC, which also plays a major role in conflict resolution as in our flanker task. It might be that P4 induces a moderate alerting state, that strengthens attention networks modulation, specifically, strengthen the ability to resolve a conflict when the attention is oriented to the target.

We did not find a significant difference in E2 concentrations in the two time-points for either group. Consequently, we cannot attribute the difference found in our study to E2 effect on attention. This null effect appears to be in agreement with some studies (e.g., Petersen et al., 2014). However, several considerations must be addressed. First, we measured E2 in the early luteal phase, several days after ovulation (18^th^ day), a time in which E2 concentrations are lower than their pre-ovulatory peak, and prior to their lower second peak in the mid-luteal phase, resulting in a more subtle differences between the groups. Second, salivatory-E2 reflects only free E2, which roughly represent circulating levels, so differences are more subtle and harder to detect. Collecting blood samples from participants, rather than saliva, would have provided an accurate measure of total E2 (i.e., bound + free), allowing the detection of subtle differences in concentrations. Finally, larger sample size may capture an effect of E2 on the three attentional networks, presuming one exists. Thus, our study cannot rule out the effect of E2 on the interaction between the three attentional networks, but only emphasize the effect of P4.

Among the participants in the present study, a significant difference in anxiety scores was found, although the sample consisted of students that did not differ in demographical factors, did not meet the criteria of anxiety, depression or premenstrual dysphoric disorders. The levels of self-reported anxiety among the NC women were higher in the early follicular phase, while in the OC group, the levels of self-reported anxiety were higher in the early luteal phase. Exploring the individual participants show that two participants from the NC group had a moderate level of anxiety in the early follicular phase, but a low level of anxiety in the early luteal phase. Since self-reports measures could be affected by situational factors, we tested mood in both phases, and excluded only individuals who met the criteria for both phases. Accordingly, these two participants were not excluded from the study. However, excluding these two participants, do not show any change in the pattern of results found.

To conclude, our study showed the effect of the menstrual cycle on the three attentional networks. Specifically, we found that in the early luteal phase of naturally cycling women alertness network was active with and without an alerting cue. The alertness network affected the other two networks, orienting and executive, influencing the ability to resolve a conflict when attention is oriented to the same location as the conflicted stimuli. We suggest that the alertness effect we found originates in the modulating effect of ovarian hormones, and specifically, to the mediating effect of progesterone on the three attentional networks; higher level of progesterone likely induced an alerting state that influenced on the ability to resolve a conflict in different orienting attention. The effect of progesterone on attention may result from the modulation of progesterone on the locus coeruleus and the anterior cingulate cortex; however, neural mechanisms should be directly studied in order to test this suggestion.

## Acknowledgments

This work was supported by the European Research Council under the European Union’s Seventh Framework Programme [FP7/2007-2013/ERC Grant Agreement no. 295664] awarded to Avishai Henik, and by the Israel Science Foundation [Grant 1799/12] in the framework of their Centers of Excellence. We wish to thank Desiree Meloul, Naama Katzin, Gal Ben Yosef, Sappir Saad, Lisa Beckmann and Dr. Daniela Aisenberg for their professional and generous help.

## Compliance with Ethical Standards

### Ethical approval

All procedures performed in studies involving human participants were in accordance with the ethical standards of the institutional and national research committee and with the 1964 Helsinki declaration and its later amendments or comparable ethical standards.

### Informed consent

Informed consent was obtained from all individual participants included in the study.

## Footnotes

1 These four exclusion criteria overlapped and some participants met two criteria.

2 When the reported menstrual cycle was 27 days, the chosen days were 3rd and 17th, and when the reported menstrual cycle was 29, the chosen days were 5th and 19th.

3 The second recruitment phase replicated the first phase’s behavioral results. Both phases, separately and together replicated the main ANT-I findings and did not differ statistically.

